# Involvement of the *doublesex* gene in body color masculinization of the blue-tailed damselfly, *Ischnura senegalensis*

**DOI:** 10.1101/2020.04.11.036715

**Authors:** Michihiko Takahashi, Genta Okude, Ryo Futahashi, Yuma Takahashi, Masakado Kawata

## Abstract

Odonata (dragonflies and damselflies) display remarkable color pattern diversity including sexual dimorphism and intrasexual polymorphism. We previously found that expression of a sex-determining transcription factor, the *doublesex* (*dsx*) gene, is associated with female color polymorphism (gynomorph for female-specific color and andromorph for male mimicking color) in the blue-tailed damselfly, *Ischnura senegalensis*. Here we investigate the function of dsx gene on thoracic coloration by electroporation-mediated RNA interference (RNAi). RNAi of the *dsx* common region changed color patterns of males and andromorphic females to patterns of gynomorphic females. Further, gynomorphic color pattern was not affected by *dsx* RNAi. The long isoform of *dsx* RNAi produced no effects, suggesting that the short isoform of *dsx* is important for body color masculinization in both males and andromorphic females. Expression pattern changes were also examined in five genes with different expression levels between sexes and female morphs. Among these genes are two melanin suppressing genes, *black* and *ebony*, that were upregulated in the *dsx*-RNAi region compared to a control region. Upregulation coincides with a gynomorphic orange color instead of the black stripe observed in males and andromorphic females. *dsx* may regulate male color differentiation by suppressing *black* and *ebony* in the thoracic region of *I. senegalensis.* Results add to the understanding of molecular mechanisms underlying the evolution of female polymorphism in Odonata.

## Introduction

Many animals and plants exhibit intraspecific color polymorphism. Color polymorphism is well studied for its ecological and evolutionary significance, including reproductive strategies, speciation, mimicry and crypsis (Gray & McKinnon, 2007). As examples, male color polymorphism of the side-blotched lizard, *Uta stansburiana*, is part of a reproductive strategy similar to the children’s game, “rock-paper-scissors” (Sinervo & Lively, 1996), and female color polymorphism seen in several butterfly species is associated with Batesian mimicry (Krushnamegh Kunte, 2009; Mallet & Joron, 1999). Many Odonata (dragonflies and damselflies) species have intraspecific body or wing color variations that are sexually dimorphic and intrasexually polymorphic (Bybee et al., 2016; Corbet 1999; Futahashi, 2016, 2017; Tillyard, 1917). Most intrasexual color polymorphisms of Odonata are female-limited and have appeared independently multiple times within the order, Odonata (Fincke, Jödicke, Paulson, & Schultz, 2005). In female-limited color polymorphism, one morph, called the andromorph, typically resembles a conspecific male, while the other morph, called the gynomorph, shows female-specific color. Among Odonata, the genus, *Ischnura*, is a model group for studies of ecology and evolution of female polymorphism. Among 75 *Ischnura* species, 29 have female-limited color polymorphism, associated with mating strategy (Rosa A. Sánchez-Guillén et al., 2018; Willink, Duryea, & Svensson, 2019). Polyandrous species often display female polymorphism, and most monandrous species are monomorphic (Robinson and Allgeyer, 1996; Sánchez-Guillén *et al*., 2018). Female-limited polymorphism in *Ischnura* is maintained by negative frequency-dependent selection and balancing selection derived from male mating harassment (Andres, Sanchez, & Cordero, 2000; Takahashi, Nagata, & Kawata, 2013; Takahashi, Kagawa, Svensson, & Kawata, 2014). In addition to body color, other phenotypic traits such as flying distance, larval duration and wing length also differ between andromorphic and gynomorphic females (van Gossum *et al.*, 2001; Abbott and Gosden, 2009). In several *Ischnura* species, female-limited color polymorphisms are controlled by an autosomal locus with two or three alleles (Cordero, 1990; Cordero & Andres, 1999; Johnson, 1964, 1966; R A Sánchez-Guillén, van Gossum, & Cordero, 2005; Takahashi et al., 2014), although little is known about the genes involved in the color polymorphisms.

Recently, a master gene that controls female-limited polymorphism was identified in several insect species (Kunte et al., 2014; Nishikawa et al., 2015; Woronik et al., 2019; Yassin, Chung, Veuille, David, & Pool, 2016; Yassin, Delaney, et al., 2016) (Kunte et al., 2014; Nishikawa et al., 2015; Yassin et al., 2016a, 2016b; Woronik et al., 2019). Among the swallowtail butterflies species in the genus, *Papilio*, a sex-determining transcription factor, the *doublesex* (*dsx)* gene, is involved in female limited Batesian mimicry (Iijima et al., 2018; Kunte et al., 2014; Nishikawa et al., 2015; Palmer & Kronforst, 2020). Expression of the *dsx* gene is also associated with female color polymorphism in the blue-tailed damselfly, *Ischnura senegalensis* (Takahashi, Takahashi, & Kawata, 2019). Adult female *I. senegalensis* display two color morphs, gynomorphic and andromorphic. The former morph is dominant.. Males and andromorphic females display a greenish color with black mid-dorsal and humeral stripes on their thorax; gynomorphic immature females display an orange color without black humeral stripes (Fig. 2). The short isoform of the *dsx* gene is expressed predominantly in males, the long isoform of *dsx* gene is expressed predominantly in females (Takahashi et al., 2019). Notably, expression of the short *dsx* isoform is higher in andromorphic females compared with gynomorphic females (Takahashi et al., 2019), suggesting that *dsx* is involved in female color polymorphism as well as sexual color dimorphism. In this study, using electroporation-mediated RNAi techniques, the effects of the *dsx* gene on adult thoracic color patterns in the blue-tailed damselfly, *I. senegalensis*, were investigated. The *dsx* gene is essential for masculinization of body color but is not involved in expression of the gynomorphic color pattern. Also, *dsx* knockdown in males and andromorphic females promoted expression of melanin and suppressed genes, *black* and *ebony*, compared to expression in gynomorphic females. The *dsx* gene may regulate masculinization of thoracic body color through the repression of *black* and *ebony* genes in *I. senegalensis*.

## Materials and Methods

### Insects

Adult female *I. senegalensis* were collected at Heiwasouzou, Makabe, Kakinohana and Shikiya on Okinawa Island in May 2017, April and October 2018, and June 2019. Mature females were placed in plastic cups (diameter 11 cm, height 4 cm) with a wet paper filter at room temperature for oviposition. Newly hatched larvae were reared together for approximately 1 month, then transferred to small plastic containers (diameter 3 cm, height 5 cm) for individual rearing. Larvae were fed with *Artemia* brine shrimp and *Tubifex* worms until early final instar. Sexes of larvae were judged by the presence of an ovipositor under the abdomen. Although color morphs of larvae cannot be determined, genetically recessive andromorphic females were expected to be obtained from eggs derived from andromorphic females. It should be noted that female morph frequency differs significantly among populations (e.g., gynomorph frequency is high (>90%) at Makabe, and low (<40%) at Kakinohana), despite small genetic distances among populations at Okinawa Island (Inomata, Hironaka, Sawada, Kuriwada, & Yama, 2015).

### Electroporation-mediated RNAi experiment

A small interfering RNA was designed using the siDirect program version 2.0 (http://sidirect2.rnai.jp/). For each gene of *I. senegalensis*, two siRNAs were designed and mixed (100 mM each). Target sequences for the *dsx* common region were 5’-AAC ATG AAT TTC GTC AAA GAT GC-3’ and 5’-ACG CAA TTC TGC AAA ATG CAA GG-3’, whereas those for *dsx* long isoform-specific region were 5’-AAC GAA AAG CAG AAA AAG GAATG-3’ and 5’-AAG TGA TAA TGC GAC GTA TAT TG-3’ (Fig. 1a). siRNA for the *dsx* short isoform-specific region was not designed because only the junction site is different. Part of long isoform would include region 2 because region 2 was shown in reverse transcription PCR of gynomorphic females (data not shown). siRNAs for the *multicopper oxidase 2* (*MCO2*) gene that is essential for melanin pigmentation (Okude, Futahashi, Kawahara-Miki, et al., 2017), were used as positive controls for targeting 5’-GAGCACTTTCCGTTATCAATATA-3’ and 5’-TCCTCTTGATGCTATCTGTAATG-3’. As a negative control, siRNA for the *enhanced green fluorescent protein* (*EGFP*) gene was used for targeting 5’-CGG CAT CAA GGT GAA CTT CAA GA-3’ (Ando & Fujiwara, 2013).

**Fig. 1.**
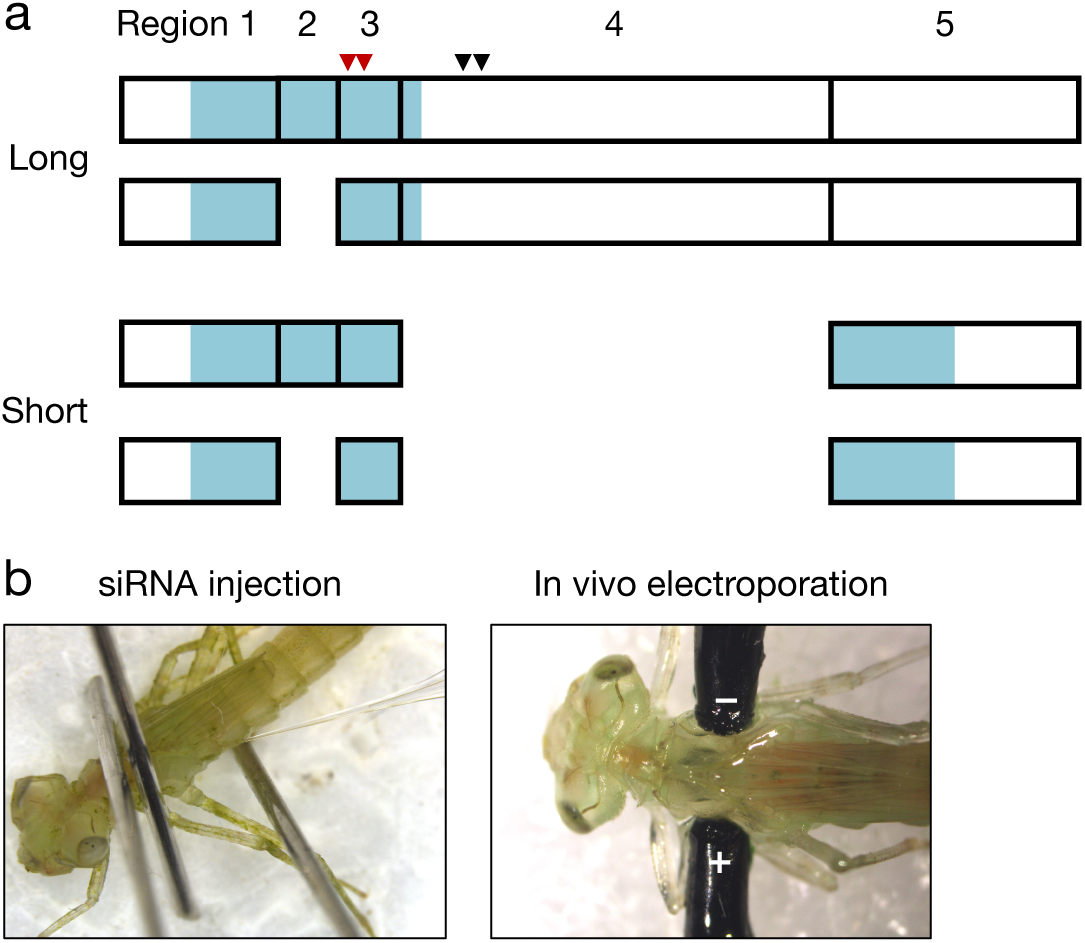
The positions of *dsx* small interfering RNAs (siRNAs)(**a**) and electroporation-mediated RNA interference (RNAi) process (**b**). **a** Summary of *dsx* isoform (modified from Takahashi et al., 2019). The red and black arrowheads indicate siRNA regions for the common region and long isoform, respectively. The blue boxes indicate open reading frames. Numbers above boxes are region identifiers. Based on RNAseq and qRT-PCR analyses, region 2 was sometimes skipped for both long and short isoforms. **b** After siRNA injection, electroporation is performed with positive and negative electrodes placed on left and right side of the thoraxes, respectively, with droplets of ultrasound gel.

Electroporation was used to introduce siRNA into larvae (Okude, Futahashi, Kawahara-Miki, et al., 2017). Early-stage, final instar larvae (stage 1 in Okude, Futahashi, Tanahashi, & Fukatsu, 2017) were anesthetized on ice for 1 min, approximately 1 μl of 100 μM siRNA solution was injected into the left side of the thorax using a glass needle and an injector (Nanoject II; Drummond Scientific Company). Immediately after injection, droplets of ultrasound gel were placed near the injection site and on the right side of the thorax opposite the injection site. Positive and negative platinum electrodes were attached to injection and opposite sites, respectively (Fig. 1b, Okude, Futahashi, Kawahara-Miki, et al., 2017). Electroporation was conducted with five or ten pulses for 25 V (each 280 ms pulses/s) using an electroporator (Cure-Gene; CellProduce). Each larva was placed on a wet paper in a plastic case individually for a day to confirm recovery from anesthesia. Injected larvae were reared at room temperature until adult emergence. Since more than half the individual larvae died before adult emergence, only adults that emerged normally were used for phenotype observation. Of note, adult phenotypes were patchy around the area where the positive electrode was placed (Fig. 2).

**Fig. 2.**
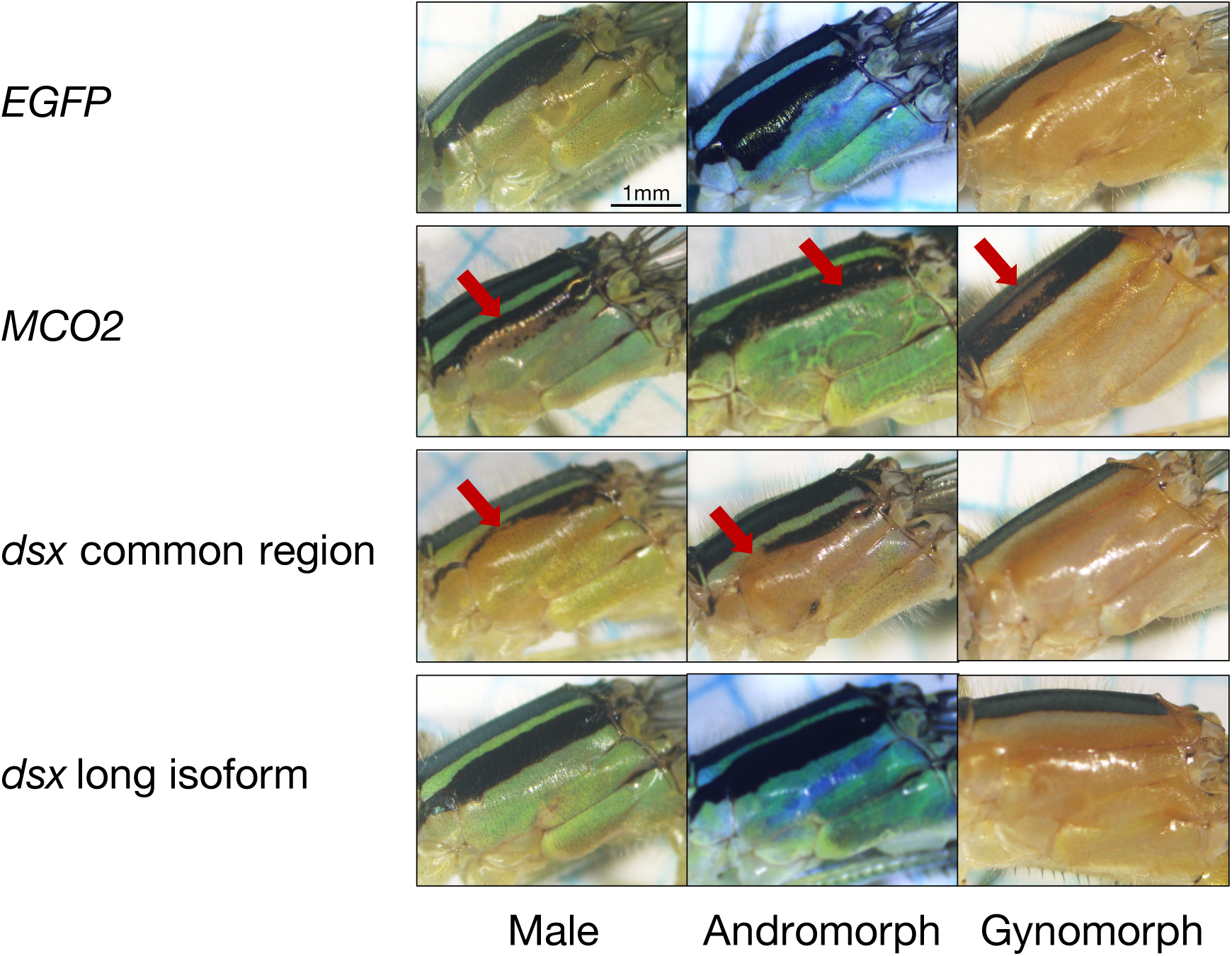
RNAi knockdown phenotypes observed in the thorax of *I. senegalensis*. Vertical axis is target genes of RNAi. No phenotypic effects in the body color of *EGFP* RNAi individuals were observed. *MCO2* RNAi individuals of all the three morphs mostly exhibited patchy unpigmented regions on the left side of the thorax (arrows) where the positive electrode was placed for electroporation. The thorax colors of males and andromorphs were changed into orange by *dsx* common region RNAi (arrows). RNAi of *dsx* long isoform did not cause any color changes in the thorax of all the three morphs.

### Quantitative PCR of adult knock-down *dsx*

To estimate RNAi efficiency of *dsx* gene electroporation, and to investigate expression of five sex and morph-associated genes (*black*, *ebony*, *chp*, *eas* and *RhoGEF*), quantitative reverse transcription polymerase chain reaction (qRT-PCR) was used. Total RNA was extracted from the left side of thorax with gynomorph-like color where siRNA was introduced and from the control region on the right side of the thorax of the same individual (N = 4 each for male and andromorphic female adults) using RNAqueous™-Micro Total RNA Isolation Kit (Thermo Fisher). cDNA was synthesized using SuperScript IV Reverse Transcriptase (Thermo Fisher) and subjected to real-time quantitative PCR using the StepOne Real-Time PCR System (Thermo Fisher) with Power SYBR Green PCR Master Mix (Applied Biosystems). Standards for each gene (*dsx* short isoform, *black*, *ebony*, *chp*, *eas* and *RhoGEF*) for qRT-PCR were generated by PCR using Tks Gflex DNA Polymerase (TaKaRa) and gene-specific primers (Table S1). The *ribosomal protein L13* (*RpL13*) gene was used as an internal standard to estimate relative mRNA expression (Table S2). The expression levels were visualized relative to maximal expression, defined as a level of 1 for each transcript. Statistically significant differences in gene expression were evaluated using the generalized linear model (GLM) assuming a gamma distribution.

## Results

### The *dsx* gene is involved in body color masculinization in *I. senegalensis*

We first investigated the effects of electroporation-mediated RNAi for *MCO2* (positive control) and *EGFP* (negative control) genes. Of fifteen individuals (four male, three andromorphic female, and eight gynomorphic female adults) that subjected to the *MCO2* RNAi and emerged normally, ten individuals (10/15 = 67%) exhibited patchy unpigmented regions on the left side of the thorax where the positive electrode was placed for electroporation (Table 1, Fig. 2, arrows). This observation is consistent with Okude, Futahashi, Kawahara-Miki, et al., (2017). After *EGFP* gene RNAi electroporation, 10 individuals (two male, one andromorphic female, and seven gynomorphic female adults) emerged normally, and no individuals showed an unpigmented area at the site of cathode placement (Table 2, Fig. 2).

**Table 1.** Summary of RNAi results for *MCO2* gene

| Morph | Emerged adults | Normal pigmentation | Partial changes of pigmentation |
| --- | --- | --- | --- |
| Males | 4 | 1 (25%) | 3 (75%) |
| Andromorphs | 3 | 1 (33%) | 2 (67%) |
| Gynomorphs | 8 | 3 (37.5%) | 5 (62.5%) |

**Table 2.**
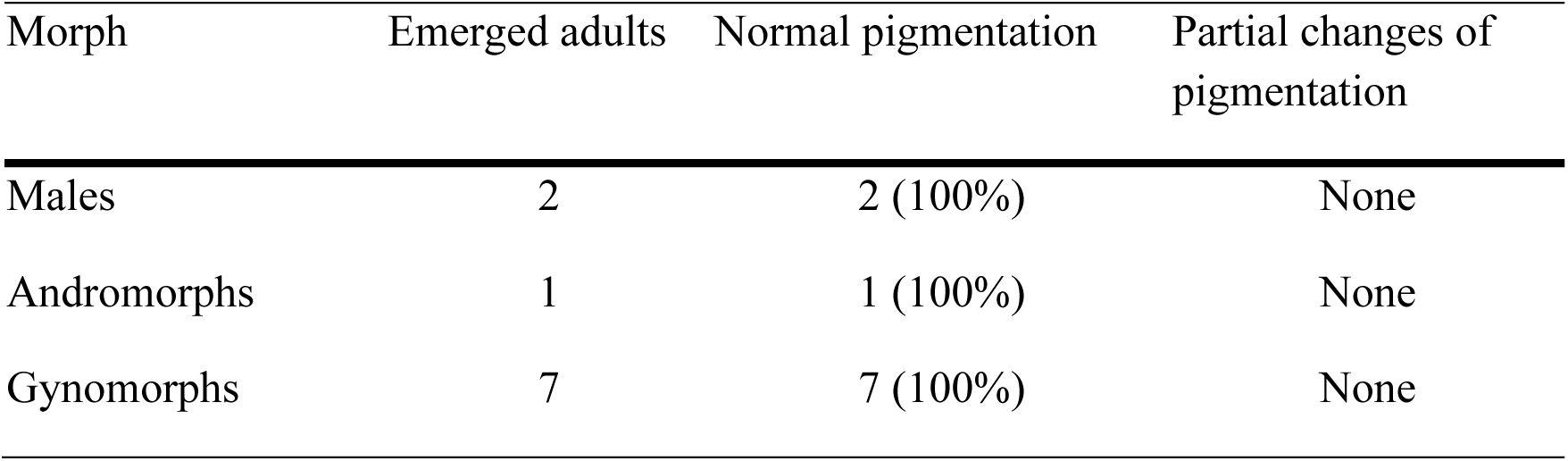
Summary of RNAi results for *EGFP* gene (negative control)

| Morph | Emerged adults | Normal pigmentation | Partial changes of pigmentation |
| --- | --- | --- | --- |
| Males | 2 | 2 (100%) | None |
| Andromorphs | 1 | 1 (100%) | None |
| Gynomorphs | 7 | 7 (100%) | None |

We next performed electroporation-mediated RNAi for the *dsx* common region (both short and long isoforms) or long isoform only. Of 41 individuals (17 male, nine andromorphic female, and 15 gynomorphic female adults) that subjected to RNAi of the *dsx* common region and emerged normally, importantly, 14 males (14/17 = 82%) and seven andromorphic females (7/9 = 78%) showed patchy orange-colored regions resembling gynomorphic females (Table 3, Fig. 2, arrows). None of the 15 gynomorphic females exhibited this color change (Table 3, Fig. 2). RNAi of *dsx* long isoform caused no noticeable effect in 31 emerged adults (nine males, four andromorphic females, and 18 gynomorphic females) (Table 4, Fig. 2). The suppression ratio for *dsx* expression (short isoform) was estimated and confirmed a significant reduction in the left side in males (*P* = 0.01963, Fig. 3); such significant reduction was not detected in andromorphic females (*P* = 0.1667, Fig. 3). Low efficiency for the reduction in target gene expression is likely due to extraction of RNA from both tissues affected by electroporation and from surrounding tissues.

**Table 3.** Summary of RNAi results for *dsx* gene (common region)

| Morph | Emerged adults | Normal pigmentation | Partial changes of pigmentation |
| --- | --- | --- | --- |
| Males | 17 | 3 (18%) | 14 (82%) |
| Andromorphs | 9 | 2 (22%) | 7 (78%) |
| Gynomorphs | 15 | 15 (100%) | None |

**Fig. 3.**
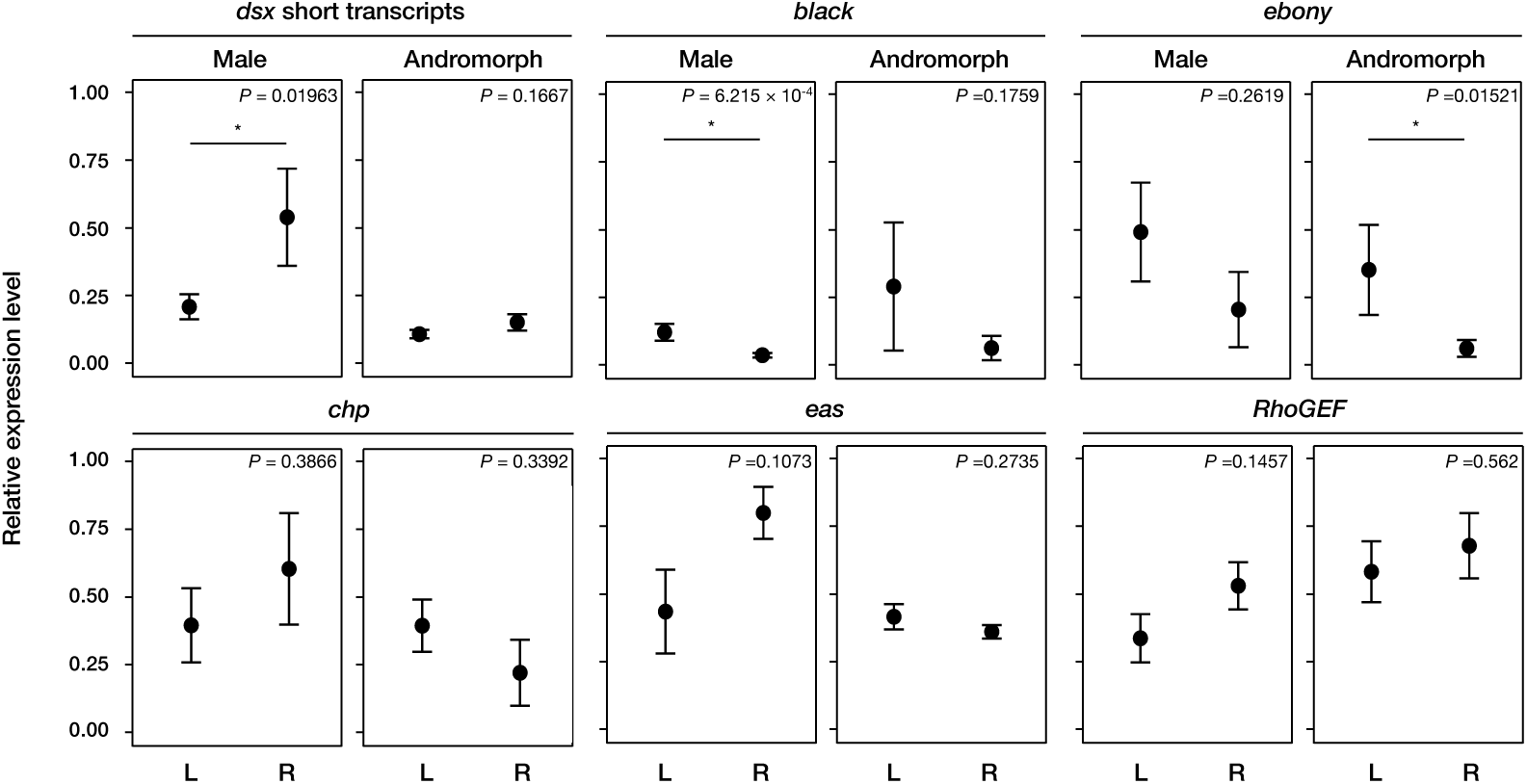
qRT-PCR analyses of *dsx* and candidates for *dsx* target genes between *dsx* (common region) RNAi region (L; left side of the thorax) and control region (R; right side of the thorax) (*N* = 4 for each male and andromorph). (a) *dsx* (short isoform), (b) *black*, (c) *ebony*, (d) *chaoptin* (*chp*), (e) *easily shocked* (*eas*) and (f) *rho guanine nucleotide exchange factor* (*RhoGEF*). Data represent mean ± S.E. Asterisks in the figure indicate significant differences (*P* < 0.05) evaluated using the Generalized Linear Model (GLM) assuming a gamma distribution.

**Table 4.** Summary of RNAi results for *dsx* gene (long isoform only)

| Morph | Emerged adults | Normal pigmentation | Partial changes of pigmentation |
| --- | --- | --- | --- |
| Males | 9 | 9 (100%) | None |
| Andromorphs | 4 | 4 (100%) | None |
| Gynomorphs | 18 | 18 (100%) | None |

### *dsx* knockdown promotes two melanin suppressing genes *black* and *ebony*

*dsx* encodes a transcription factor that regulates the expression of several downstream genes (Chatterjee, Uppendahl, Chowdhury, Ip, & Siegal, 2011; Matson & Zarkower, 2012). In addition to the *dsx* gene, *black*, *ebony*, *chaoptin* (*chp*), *easily shocked* (*eas*), and *rho guanine nucleotide exchange factor* (*RhoGEF*) genes are also differentially expressed in female morphs and between sexes (Takahashi et al., 2019). These genes represent candidate *dsx* target genes. Expression of these five genes between *dsx*-RNAi and control regions was evaluated. Expression of two melanin suppressing genes, *black* in males and *ebony* in andromorphic females, were higher on the left side (Fig. 3, asterisks *P* < 0.05) and resembled expression in gynomorphic females. No significant differences in expression were detected for the remaining three genes, *chp*, *eas* and *RhoGEF* (Fig. 3).

## Discussion

*dsx* is involved in male-type color pattern formation, but not in female-specific gynomorphic coloration in *I. senegalensis*. The male *dsx* isoform is essential for male differentiation in most insects, while the female *dsx* isoform is involved in female differentiation only in holometabolous insects (Wexler et al., 2019; Zhou, Whitworth, Pozmanter, Neville, & Doren, 2018). Our results are consistent with previous work on male differentiation. Like *Ischnura* damselflies, some butterflies display female color polymorphism, in which one morph resembles male coloration (Kunte et al., 2014; Palmer & Kronforst, 2020). Functional analysis suggests that changes in total expression level and protein sequences, but not isoform patterns, are important for female polymorphisms in butterflies (Nishikawa et al., 2015). These findings contrast with the current results for *Ischnura* damselflies.

Based on qRT-PCR analyses, *black* and *ebony* were identified as *dsx* target gene candidates. *black* and *ebony* encode aspartate decarboxylase and *N*-β-alanyldopamine (NBAD) synthase, respectively. Both enzymes are essential for the synthesis of NBAD that is involved in hardening of the cuticle, and for reddish and yellowish coloration (Arakane, Noh, Asano, & Kramer, 2016; Futahashi & Fujiwara, 2005; Liu, Lemonds, Marden, & Popadić, 2016; Wittkopp, Carroll, & Kopp, 2003). Based on phenotypes of other insects, upregulation of *black* and *ebony* is consistent with a color change from black humeral stripes to orange. These results strongly suggest that *dsx* suppresses *black* and *ebony* in males and andromorphic females, and induces a change from a female to a male color pattern. Elucidation of the regulatory mechanism of *dsx* splicing patterns may clarify the evolution of female color polymorphism as well as sexual color dimorphism in Odonata.

## Acknowledsments

We thanks Okinawa Prefectural Agricultural Research Center for collecting samples, Shinichiro Maruyama for advices of experiments and Koji Tamura for access to his real-time quantitative PCR equipment. This study was partially supported by JSPS Research Fellowship for Young Scientists and

## Data Accessibility

Data of RNAi adult phenotypes will be available in Dryad. The expression levels of *dsx* and *dsx* target genes from real-time quantitative PCR are listed in Supplementary Table S2.

## Author Contributions

M. T., Y. T. and M.K. designed research. M. T., G. O. and R. F. conducted RNAi experiment. M. T. conducted real-time quantitative PCR experiment and statistical analysis. M. T. wrote the first draft manuscript, and all authors contributed to the improvement of the manuscript.

